# The antidepressant effects of asperosaponin VI are mediated by the suppression of microglial activation and reduction of TLR4/NF-ĸB induced IDO expression

**DOI:** 10.1101/2020.03.15.992529

**Authors:** Jinqiang Zhang, Saini Yi, Yahui Li, Chenghong Xiao, Chan Liu, Weike Jiang, Changgui Yang, Tao Zhou

**Author notes:** Corresponding author: Prof. Tao Zhou, Guizhou University of Traditional Chinese Medicine, Guiyang 550025, China,.

## Abstract

**Aim:** Indoleamine 2, 3-dioxygenase (IDO) is responsible for the progression of the kynurenine pathway, which has been implicated in the pathophysiology of inflammation-induced depression. It has been reported that asperosaponin VI (ASA VI) could play a neuroprotective role through anti-inflammatory and antioxidant. In this study, we examined the antidepressant effect of ASA VI in LPS-treated mice and further explored its molecular mechanism by insight into the microglial kynurenine pathway.

**Methods:** To produce the model, lipopolysaccharide (LPS) (0.83 mg/kg) was administered intraperitoneally to mice. The mice received ASA VI (10 mg/kg, 20mg/kg, 40mg/kg and 80mg/kg, i.p.) thirty minutes prior to LPS injection. Depressive-like behaviors were evaluated based on the duration of immobility in the forced swim test. Microglial activation and inflammatory cytokines were detected by immunohistochemistry, real-time PCR and ELISA. The TLR4/NF-ĸB signaling pathway and the expression of IDO, GluA2, and CamKIIβ were measured by western blotting.

**Results:** ASA VI demonstrated significant antidepressant activity in the presence of LPS on immobility and latency times in the forced swim test. The LPS-induced activation of microglia and inflammatory response were inhabited by ASA VI in a dose-dependent manner. TLR4/NF-κB signaling pathway also was suppressed by ASA VI in the hippocampus and prefrontal cortex of LPS-treated mice. Furthermore, ASA VI inhibited the increase in IDO protein expression and normalized the aberrant glutamate transmission in the hippocampus and prefrontal cortex as a result of LPS administration.

**Conclusion:** Our results propose a promising antidepressant effect for ASA VI possibly through the downregulation of IDO expression and normalization of the aberrant glutamate transmission. This remedying effect of ASA VI could be attributed to suppress microglia-mediated neuroinflammatory response via inhibiting the TLR4/NF-κB signaling pathway.

## 1. Introduction

Depression is highly prevalent worldwide and a leading cause of disability. However, the medications currently available to treat depression fail to adequately relieve depressive symptoms for a large number of patients. Inflammation-induced depression is a debilitating psychiatric disorder that is caused by abnormal tryptophan metabolism in the central nervous system (CNS)(Dantzer et al. 2008; Kohler et al. 2016). This type of depression is refractory to conventional medications such as selective serotonin reuptake inhibitors (SSRIs)(Al-Harbi 2012; Hashmi et al. 2013). Research into the aberrant overactivation of the kynurenine pathway and the production of various active metabolites has brought new insight into the progression of depression (Ogyu et al. 2018; Oxenkrug 2010; Savitz 2017). In the non-challenged brain, kynurenine (KYN) is predominantly metabolized to kynurenic acid (KYNA) by astrocytes(Schwarcz et al. 2012). However, disruption of neuroendocrine or neuroinflammatory balance, which involves activation of microglia, KYN is metabolized to 3-hydroxykynurenine (3HK) in microglia (Schwarcz et al. 2012; Schwarcz and Stone 2017). The 3HK is further metabolized into quinolinic acid (QUIN) which is the agonist of the N-methyl-D-aspartate (NMDA) receptor that leads to neuronal dysfunction and aberrant glutamate transmission (Duman and Aghajanian 2012; Kubicova et al. 2013).

Microglia, the principal immune cells in the brain, plays a central role in immune surveillance and inflammatory-related neuropathology (Ginhoux et al. 2010; Hanisch and Kettenmann 2007; Yirmiya et al. 2015). The dysfunction of microglia has been associated with depression (Brites and Fernandes 2015; Frick et al. 2013; Yirmiya et al. 2015). The cascade of microglial activation could promote the synthesis and secretion of a large number of inflammatory mediators, resulting in neuroinflammatory response (Zhang et al. 2017; Zhang et al. 2016). Inflammatory cytokines such as IL-1β and TNF-α have been shown to enhance this pathway in microglia by potentiating the expression of the kynurenine pathway’s main controller enzyme, indoleamine 2, 3-dioxygenase (IDO), through STAT1 activation (Hemmati et al. 2019). In a similar manner, lipopolysaccharides (LPS) upregulate IDO levels through activating the TLR4/NF-ĸB signaling pathway and induce depressive-like behaviors (O’Connor et al. 2009b).

A growing body of research suggests that natural products have a good antidepressant effect by inhibiting microglia-mediated neuroinflammation (Lee et al. 2013; Zhang et al. 2017). The active ingredients of traditional Chinese medicine have great potential in anti-depression research. Asperosaponin VI (ASA VI), a natural compound isolated from the well-known traditional Chinese herb Radix Dipsaci, has an important role in promoting osteoblast formation(Ke et al. 2016). Studies have also shown that ASA VI has a neuroprotective effect through anti-inflammatory and antioxidant effects (Wang et al. 2018; Yang et al. 2019; Yu et al. 2012). In this study, we explored the antidepressant effect of ASA VI in LPS-treated mice, and the molecular mechanism was further explored by detecting the activation of microglia, TLR4/NF-κB pathway, IDO protein expression, and GluA2 and CamKIIβ.

## 2. Materials and methods

### 2.1 Animals

Male C57BL/6 mice (8 weeks old) were purchased from the Biotechnology Centre (Chang Sha, China). All the animals were raised in a single cage in a suitable environment of 25 ± 1 °C and 65% humidity, with free drinking water and activities. All animal experiments were carried out by the China code for the care and use of animals for scientific purposes at Guizhou University of Traditional Chinese Medicine, Guiyang, China, with local animal ethics committee approval.

### 2.2 LPS administration and pharmacological intervention

The asperosaponin VI (ASA VI) standard (purity = 99.92%) was purchased from Chengdu Alfa Biotechnology Co., Ltd. The asperosaponin VI was dissolved in saline. To produce the model, lipopolysaccharide (LPS) (0.83 mg/kg) was administered intraperitoneally to mice. The control mice were administered with equivoluminal saline. Both control mice and model mice were received ASA VI (10 mg/kg, 20mg/kg, 40mg/kg and 80mg/kg, i.p.) thirty minutes prior to LPS injection. Following the behavioral analysis, immunocytochemistry, RT-PCR analysis, ELISA and western blot analysis were performed.

### 2.3 Open field test (OFT)

The open field test was used to ensure that alterations in FST immobility duration were not related to changes in locomotor function. For this test, each mouse was placed in an acrylic plastic box (50 × 50 × 30 cm). The a locomotor activity, the time in center and travel distance during a 5-min period were measured by OFT 100 software (Taimen, Chengdu, China).

### 2.4 Forced swim test (FST)

Each mouse was placed in a cylinder containing 15 cm water for 15 min. After 24 h, the animals were placed again in the cylinder and the duration of immobility (including movements to keep afloat, but not active swimming) was scored for a 6 min period.

### 2.5 Immunocytochemistry and image analysis

Mice were anesthetized with 1% pentobarbital (50 mg. Kg^-1^. i.p.), perfused with 4% paraformaldehyde (0.1 M phosphate buffer configuration, PBS) for 10 minutes, the whole brain of mice was isolated and put into 4% paraformaldehyde for 48 hours. Then dehydrated in 30% sucrose for 48 hours, frozen section was 30μm. Brain slices were placed in 24-well plates, wash with PBS for 3 times, 5 min/ time, permeate with 0.5% Triton x-100 for 15 min, then wash with PBS for 3 times again, 5 min/ time. The brain slices were incubated for 2 h at room temperature in a blocking solution (10% Donkey Serum prepared with PBS) and incubated for 24 h at 4 °C in the solution, including the primary antibodies (mouse anti-Iba1 antibody, Abcam, 1:400; rabbit anti-iNOS antibody, Abcam, 1:50; mouse anti-GFAP antibody, Cell signaling, 1:400; rabbit anti-NeuN antibody, Cell signaling, 1:400). After incubation, the brain slices were incubated with the secondary antibodies (anti-rabbit IgG-conjugated Alexa Fluorochrome or anti-mouse IgG conjugated Alexa Fluorochrome, Invitrogen; 1:500) for 2 h at room temperature.

Microglia and astrocyte were imaged using fluorescence microscopy (Olympus BX51). Image analysis was used to quantify the area, perimeter and fluorescence intensity of microglia. The images of microglia were imported to Image J software (version 1.45 J) and used to determine a threshold for positive staining while excluding background staining. The amount, average percent area, fluorescence intensity of the positive threshold for all representative pictures are reported.

### 2.6 Quantitative PCR

Mice were sacrificed under aseptic conditions, and the hippocampus and cortex were isolated and placed in 1.5 mL aseptic centrifuge tubes, total RNA was extracted by Trizol (TaKaRa, Japan), use the reverse transcription kit(Invitrogen Life Technologies, USA)to get the cDNA in strict accordance with the steps in the experimental instructions. The RT-PCR reaction mixture contains 1 μL of template cDNA, 5 μL MasterMix and 1 μL primer (Sangon Biotech, Sichuan, China), add DEPC water to a total reaction volume of 10 μL. After mixing, put it into the 7500 Real-Time PCR System (Applied Biosystems, USA). The internal reference gene is β-actin and the expression of related genes are calculated according to the method of -ΔΔ Ct. Primer sequences have been listed in Table S1.

### 2.7 Enzyme-linked immunosorbent assay (ELISA)

Mice were sacrificed under aseptic conditions, and the hippocampus and cortex of the mice were placed in a 1.5 mL centrifuge tube. After homogenization, the supernatant was centrifuged for testing the secretion of inflammatory cytokines. BCA (Sangon Biotech, Sichuan, China) was used to determine the concentration of total protein, diluted to the same concentration, and then operated in strict accordance with the instructions of the ELISA Kit (BOSTER, Wuhan, China) to determine the levels of IL-1β, and TNF-α.

### 2.8 Western blot analysis

Mice were sacrificed under aseptic conditions, and the hippocampus and cortex of the mice were placed in a 1.5 mL centrifuge tube, RIAP lysis buffer (Solarbio, China) was used for lysis. The lysis solution was centrifuged at 1000×g for 30 min. The concentration was measured by the BCA method. Equal amount of protein was resolved using 12% SDS polyacrylamide gel. The proteins were transferred onto PVDF membranes. The membranes were blocked with 5% skimmed milk and incubated with primary antibodies (rabbit anti-PPAR-γ antibody, Cell Signaling Technology, 1:1000) and HRP-conjugated goat anti-rabbit IgG. The proteins were tested using the chemiluminescence detection system (Amersham, Berkshire, UK). Finally, the bands were analyzed using Image J software (version 1.45 J).

### 2.9 Statistic analysis

The statistical analyses were performed using SPSS 17.0 software (SPSS Inc., Chicago, USA). Differences between the mean values were evaluated by a paired Student’s t-test for independent samples, one-way or two-way analysis of variance (ANOVA) followed by a protected Fisher’s least significance difference (LSD) test for post hoc comparisons of multiple groups. A P value of 0.05 or less was considered indicative of statistical significance. Each sample was repeated 3 times for qPCR, ELISA and western blot, 5 immunofluorescence images of each sample were used to image analysis. The mean value of the parallel repeated data was used for significance statistical analysis.

## 3. Results

### 3.1 ASA VI alleviates LPS-induce depressive-like behaviors

Motor function, immobility, and latency times were compared between study groups of 8–15 mice. In the OFT, changes in mobility and stationary time and moved distance were not significant following ASA VI pretreatment regardless of LPS or saline administration as determined by two-way ANOVA (Fig. 1B and 1C). The changes in the time in center were not significant following ASA VI pretreatment in control mice when compared with saline-treated mice. LPS treatment markedly decreased the time in the center of mice in the OFT. While ASA VI pretreatment reversed the LPS-induced increase in the time in center in a dose-dependent manner. All of the 20 mg/kg, 40 mg/kg and 80 mg/kg of ASA VI except the dose of 10 mg/kg significantly increased the time in center when compared with LPS-treated mice without pharmacological intervention (Fig. 1D).

**Fig. 1.**
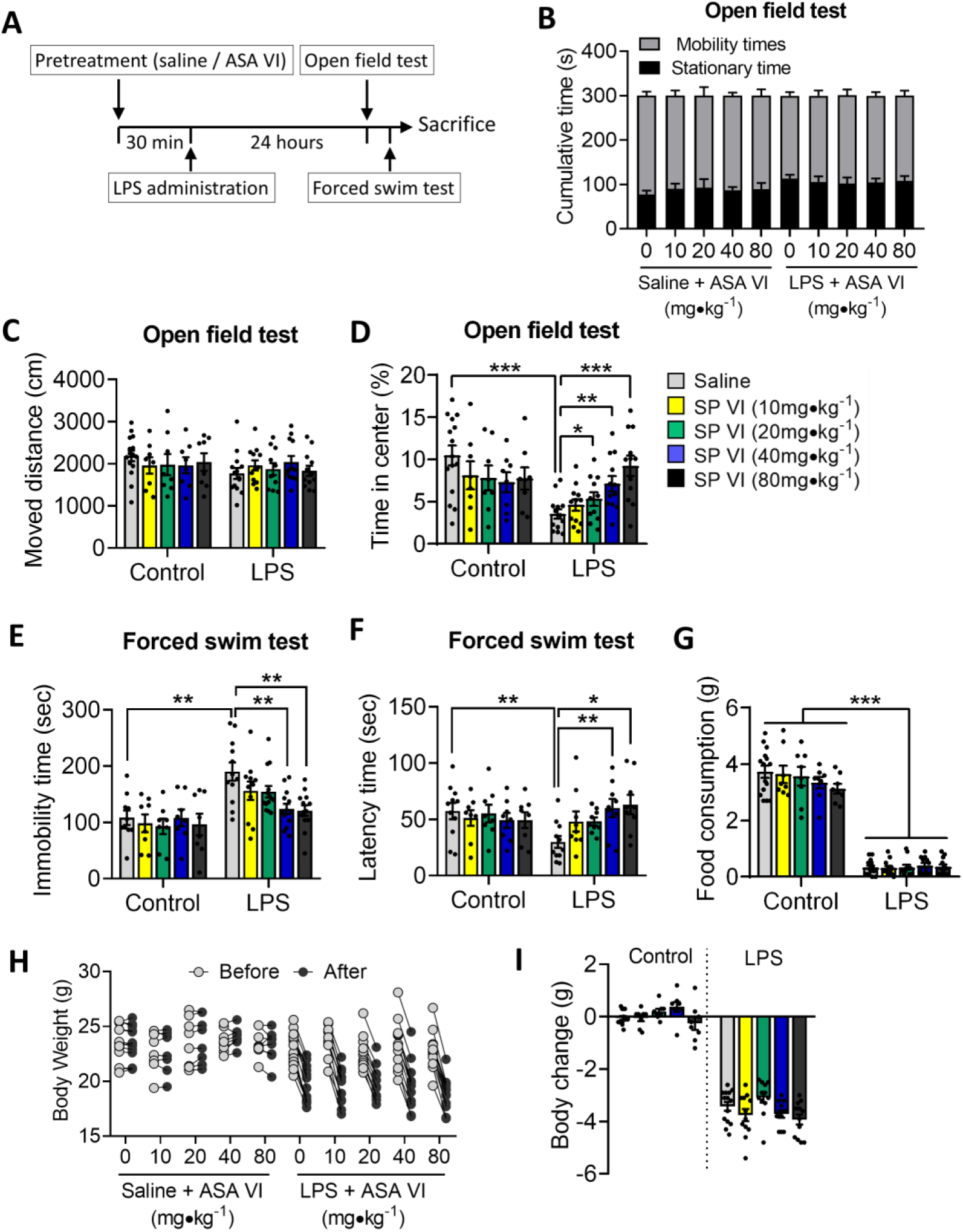
Effects of ASA VI on LPS-induce depressive-like behaviors. **A**: The timeline of our experimental process. **B-D**: Changes in mobility and stationary time, moved distance and time in center of mice in open field test (OPT) following ASA VI pretreatment before LPS or saline administration. **E** and **F**: Changes in immobility and latency times of mice in forced swim test (FST) following ASA VI pretreatment before LPS or saline administration. **G**: Changes in food consumption of mice during 24 h following ASA VI pretreatment before LPS or saline administration. **H** and **I**: Changes in body weight of mice during 24 h following ASA VI pretreatment before LPS or saline administration. Data of figure 1B are mean ± SEM, data of figure 1C-1I is displayed individually (n =8-15 per group), * P < 0.05, ** P < 0.01, *** P < 0.005 (two-way ANOVA with LSD test).

In the FST, changes in immobility and latency times were not significant following ASA VI pretreatment in control mice when compared with saline-treated mice. (Fig. 1E and 1F). LPS treatment markedly increased the immobility time and decreased the latency times in the TST. While ASA VI pretreatment reversed the LPS-induced increase in the immobility time and decrease in latency times in a dose-dependent manner. The 40 mg/kg and 80 mg/kg but not 10 mg/kg and 20 mg/kg of ASA VI significantly decreased in the immobility time and increased latency times in FST when compared with LPS-treated mice without pharmacological intervention (Fig. 1E and 1F).

The food consumption and body weight of LPS- or saline-treated mice were examined when pretreated with 10 mg/kg, 20 mg/kg, 40 mg/kg and 80 mg/kg of ASA VI. The results showed that several doses of ASA VI did not reserve the LPS-induced decrease in food consumption and body weight (Fig. 1G – 1I). It is worth noting that 80 mg/kg of ASA VI generally reduced the weight of control mice.

### 3.2 Antidepressant effect of ASA VI is potentially mediated by inhibiting LPS-induced activation of microglia and neuroinflammation

In the brain, glia could respond positively to LPS stimulation. We examined the effects of ASA VI on the astrocyte and microglia in the hippocampus and PFC of LPS-treated mice. The results showed that the changes in GFAP mRNA expression and area of GFAP^+^ staining were not significant following ASA VI pretreatment regardless of LPS or saline administration as determined by two-way ANOVA (Fig. 2A – 2C). LPS administration results in the increase of microglial markers (Iba1 and CD11b) expression and change of the microglial morphology, including the increased area of microglia and decreased number and length of branches in the hippocampus and PFC (Fig. 2). The ASA VI inhibited LPS-induced activation of microglia in a dose-dependent manner. When the mice were pretreated with ASA VI (40mg/kg, 80mg/kg) before LPS administration, the mRNA expression of microglial markers (Iba1 and CD11b) were significantly inhibited in hippocampus and PFC (Fig. 2D – 2G). The 10 mg/kg ASA VI were not significantly affected the mRNA expression of microglial markers. The effects of 20 mg/kg ASA VI on the mRNA expression of microglial markers are wobbly in the hippocampus and PFC of LPS-treated mice (Fig. 2D – 2G).

**Fig. 2.**
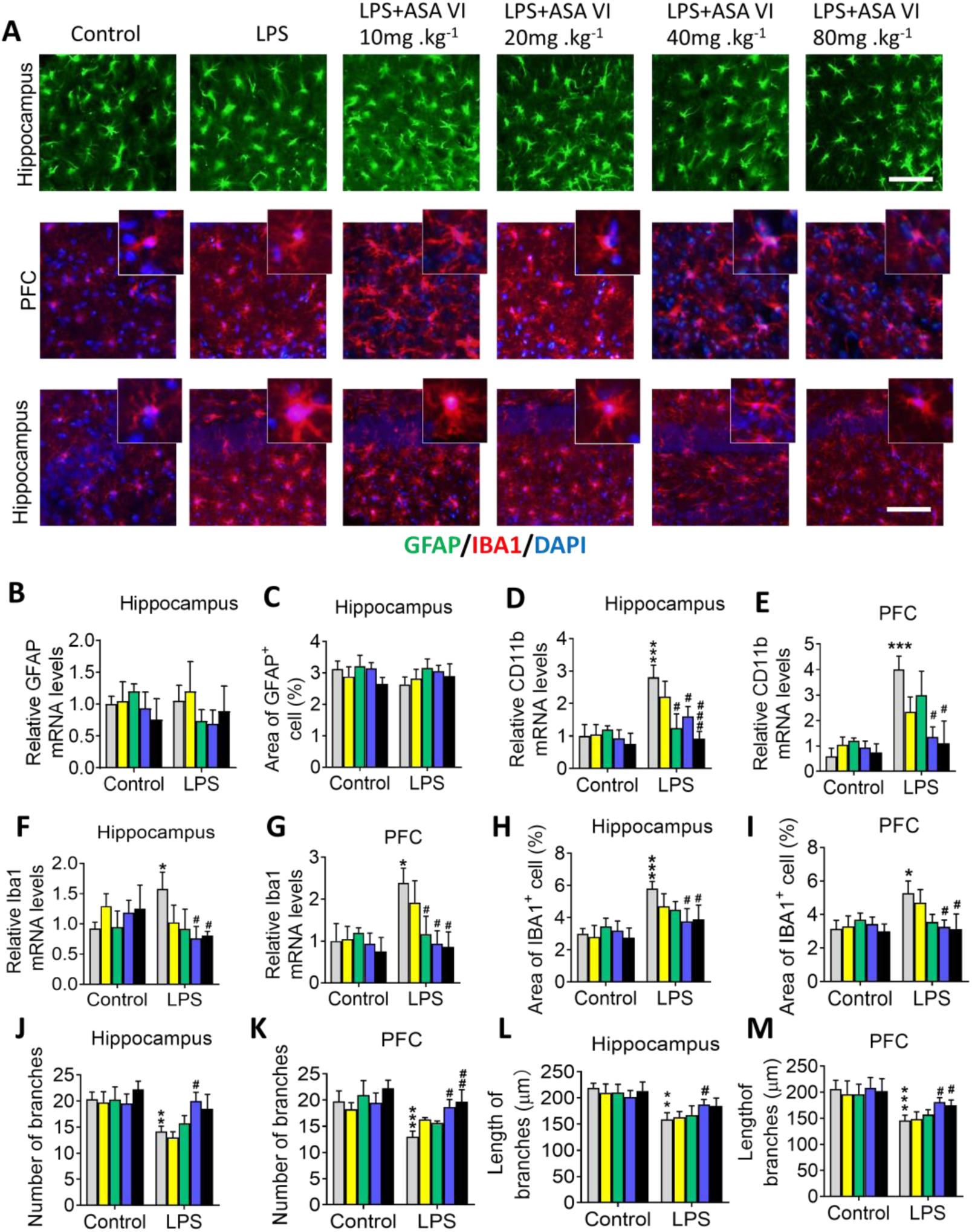
Effects of ASA VI on LPS-induced activation of microglia in hippocampus and prefrontal cortex. **A**: Immunohistochemistry detects the changes of astrocyte (GFAP, green) and microglia (IBA1, red) in hippocampus and prefrontal cortex of mice following ASA VI pretreatment before LPS or saline administration. Scale bar, 50μm. **B**: Quantitative PCR detects the mRNA expression of GFAP in hippocampus. Data are showed the fold change relative to control group. **C**: Quantification of the area of GFAP^+^ cells in hippocampus of mice. **D-G**: Quantitative PCR detects the mRNA expression of CD11b and Iba1 in hippocampus and prefrontal cortex. Data are showed the fold change relative to control group. **H-M**: Quantification of the area, number and length of branches of IBA1^+^ cell in hippocampus and prefrontal cortex. Data are mean ± SEM (n = 4-6 per group), each sample was repeated 3 times for q-PCR, 5 immunofluorescence images of each simple were used to analysis. *** P < 0.005 when compared with control group, ^#^ P < 0.05, ^##^ P < 0.01, ^###^ P < 0.005 when compared with LPS group (two-way ANOVA with LSD test).

The LPS-induced increase in the area of IBA1^+^ cells in the hippocampus and PFC is suppressed by 40mg/kg and 80mg/kg of ASA VI. The 10 mg/kg and 20 mg/kg of ASA VI were not significantly affected the area of IBA1^+^ cells in the hippocampus and PFC of LPS-treated mice (Fig. 2H and 2I). The LPS-induced decrease in number and length of branches of microglia in the hippocampus and PFC is reserved by 40mg/kg of ASA VI. The 80 mg/kg of ASA VI only increased the number and length of branches of microglia in PFC (Fig 2J – 2M).

The expression of inflammatory cytokines is usually synchronized with changes in microglial morphology. We also examined the effects of ASA VI on the expression of proinflammatory cytokines in the hippocampus and PFC of LPS-treated mice. The results showed that the changes in mRNA expression (IL-1β, TNF-α, IL-6, and iNOS) and protein expression (IL-1β and TNF-α) of inflammatory cytokines were not significant following ASA VI pretreatment in hippocampus and PFC of control mice as determined by two-way ANOVA (Fig. 3A and 3B). LPS administration results in an increase of IL-1β, TNF-α, IL-6 and iNOS expression in the hippocampus and PFC (Fig. 3). ASA VI decreased the expression of IL-1β, TNF-α, IL-6 and iNOS in both the hippocampus and PFC of LPS-treated mice in a dose-dependent manner (Fig. 3A). The expression of IL-1β and TNF-α were further validated at the protein level by ELISA. The results were consistent with those of mRNA expression (Fig. 3B). The result from immunohistochemistry showed that LPS-induces inducible nitric oxide synthase (iNOS) was mainly expressed in microglia (Fig. 3C).

**Fig. 3.**
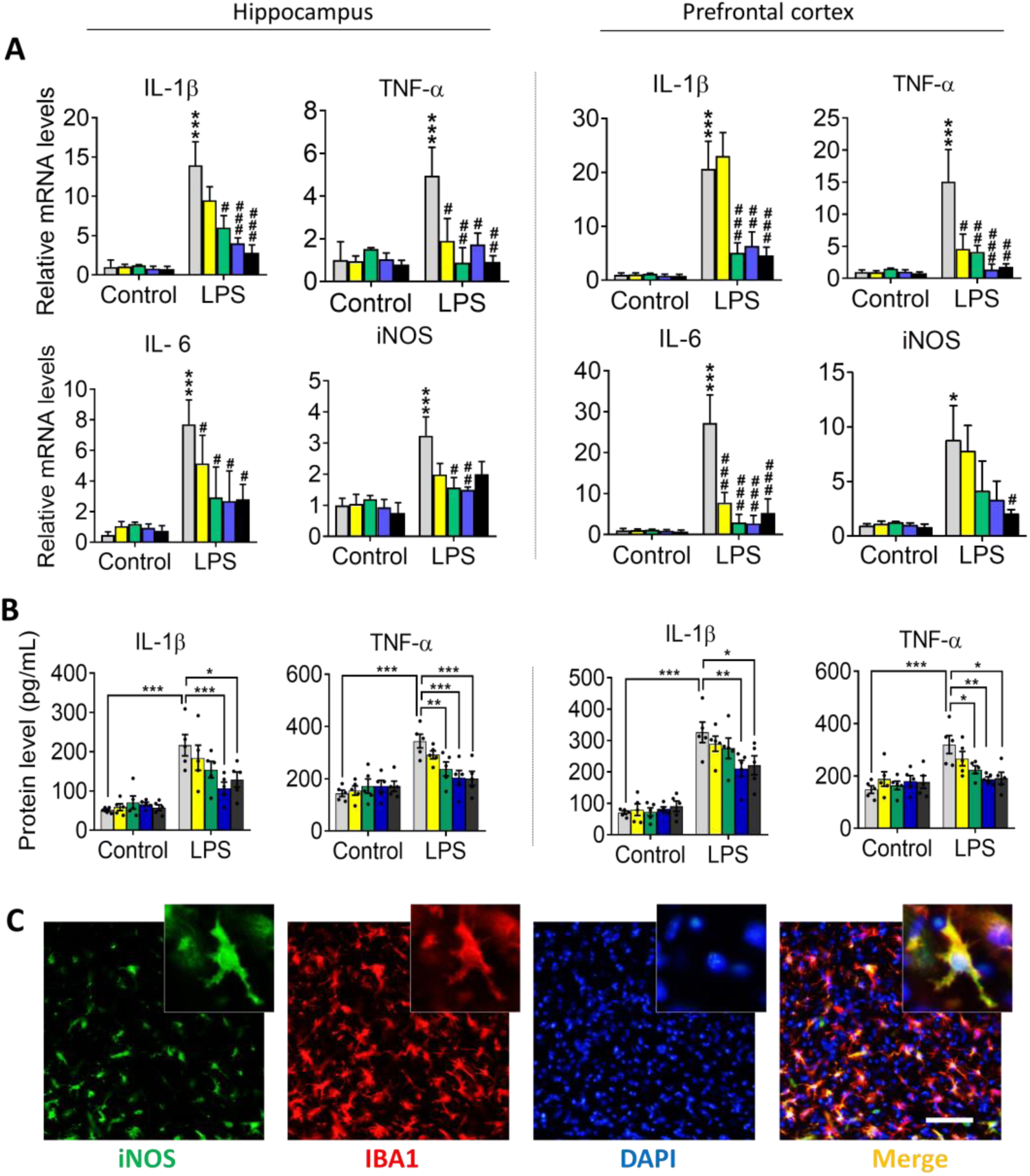
Effects of ASA VI on neuroinflammatoy response in hippocampus and prefrontal cortex of LPS-treated mice. **A**: Quantitative PCR detects the mRNA expression of IL-1β, TNF-α, IL-6 and iNOS in hippocampus and prefrontal cortex. Data are showed the fold change relative to control group. Data are mean ± SEM (n = 4-6 per group), each sample was repeated 3 times. * P < 0.05, *** P < 0.005 when compared with control group, ^#^ P < 0.05, ^##^ P < 0.01, ^###^ P < 0.005 when compared with LPS group (two-way ANOVA with LSD test). **B**: ELISA detects the protein expression of IL-1β and TNF-α in hippocampus and prefrontal cortex. Data were displayed individually (n =5 per group), each sample was repeated 3 times. * P < 0.05, ** P < 0.01, *** P < 0.005 (two-way ANOVA with LSD test). **C**: Immunohistochemistry detects the sources of iNOS (green). iNOS (green) was located in IBA1^+^ cell (red) in prefrontal cortex of mice following LPS administration. Scale bar, 50μm.

### 3.3 ASA VI suppressed microglia-mediated neuroinflammation by inhibiting the TLR4/NF-κB signaling pathway

To further explore the molecular mechanism of the anti-inflammatory activity of ASA VI, we examined the total NF-κB and phosphorylated NF-κB (pNF-κB) by western blotting in hippocampus and PFC of mice received LPS administration and ASA VI pretreatment. The results showed that LPS administration promoted the phosphorylation of NF-κB in both the hippocampus and PFC when compared with control mice (Fig. 4A – 4D). ASA VI inhibited the phosphorylation of NF-κB in a dose-dependent manner. The 40 mg/kg and 80 mg/kg of ASA VI significantly inhibited LPS-induced phosphorylation of NF-κB in both the hippocampus and PFC (Fig. 4A – 4D).

**Fig. 4.**
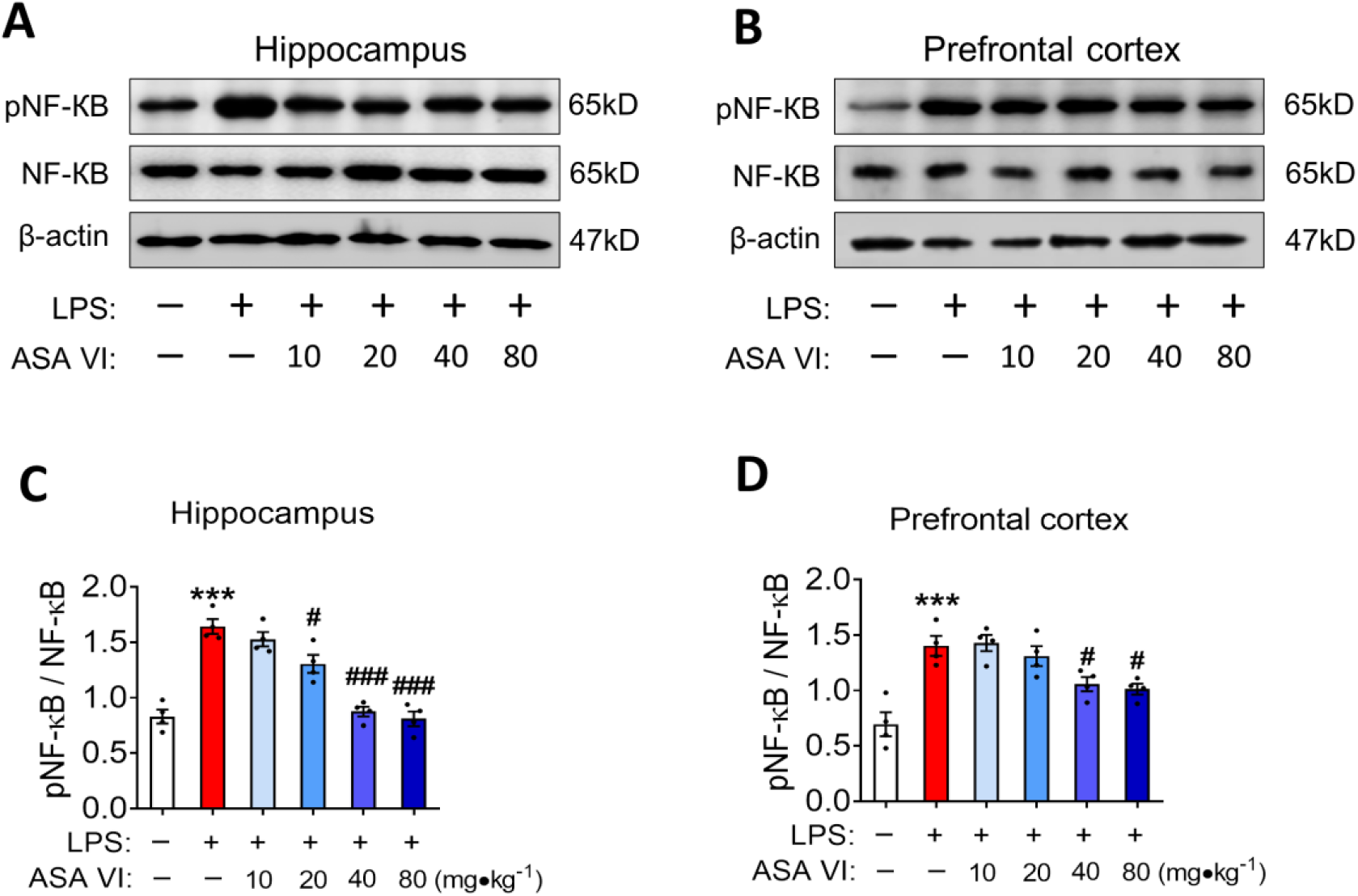
Effects of ASA VI on phosphorylation of NF-ĸB in hippocampus and prefrontal cortex of LPS-treated mice. **A** and **B**: Western blotting detects NF-ĸB and pNF-ĸB in hippocampus and prefrontal cortex of mice following ASA VI pretreatment before LPS administration. **C** and **D**: Quantification of the phosphorylation level of NF-ĸB in hippocampus and prefrontal cortex. NF-ĸB was normalized β-actin, and pNF-ĸB was normalized total NF-ĸB. Data were displayed individually (n =3-4 per group), each sample was repeated 3 times. *** P < 0.005 when compared with control group, ^#^ P < 0.05, ^###^ P < 0.005 when compared with LPS group (one-way ANOVA with LSD test).

### 3.4 IDO expression is downregulated in ASA VI-treated mice in a dose-dependent manner

Based on the effect of microglia-mediated inflammatory response on tryptophan metabolism, we examined the expression of key enzymes (IDO) in the kynurenine pathway. The protein expression of IDO1 was detected by western blotting in the hippocampus and PFC of mice received LPS administration and ASA VI pretreatment. The results showed that LPS administration increased the expression of IDO1 in both the hippocampus and PFC when compared with control mice (Fig. 5A – 5D). When the mice were pretreated with ASA VI (40 mg/kg, 80 mg/kg) before LPS administration, the expression of IDO1 in the hippocampus and PFC were inhibited when compared with the mice treated only LPS (Fig. 5A – 5D).

**Fig. 5.**
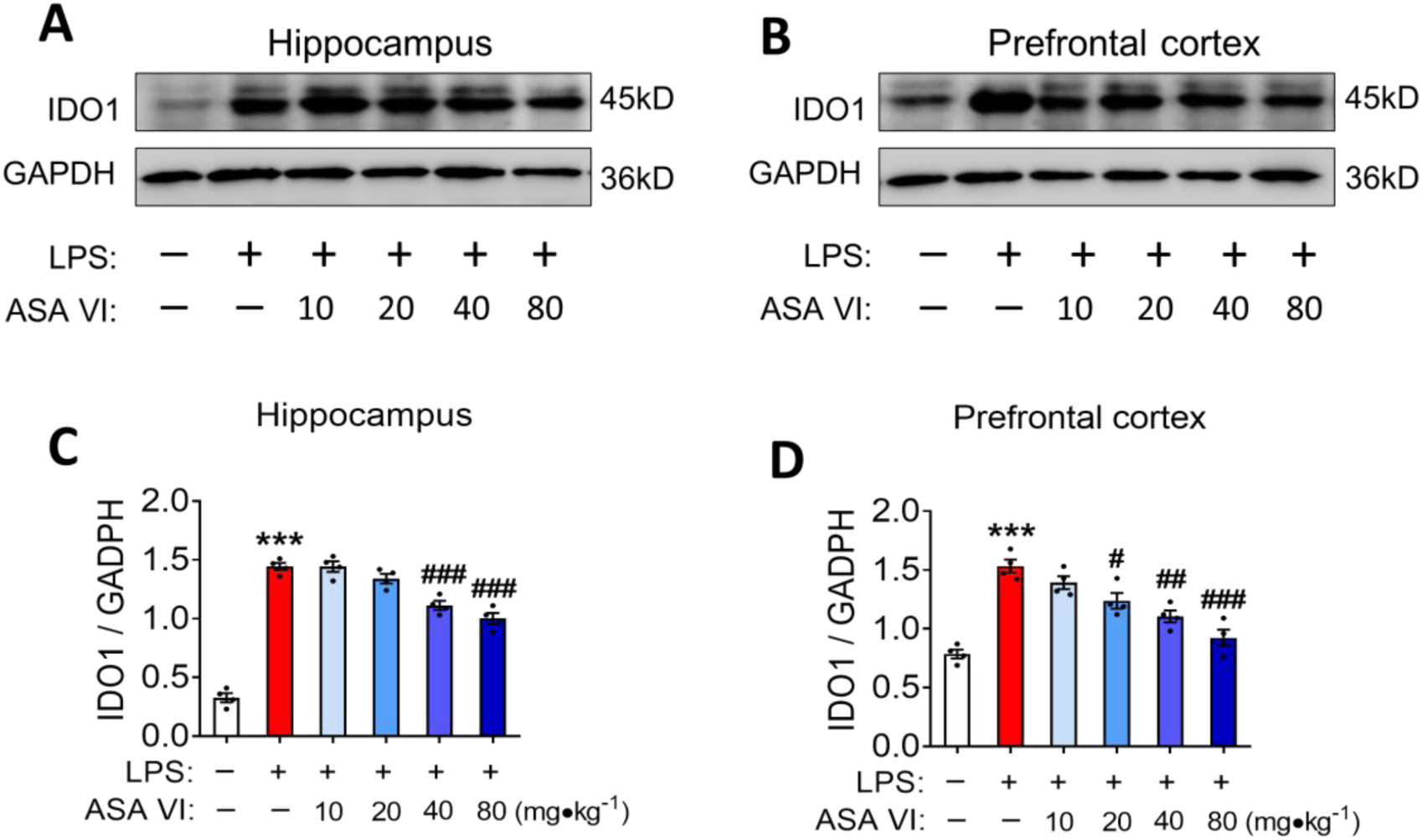
Effects of ASA VI on IDO expression in hippocampus and prefrontal cortex of LPS-treated mice. **A** and **B**: Western blotting detects IDO1 in hippocampus and prefrontal cortex of mice following ASA VI pretreatment before LPS administration. **C** and **D**: Quantification of the protein level of IDO1 in hippocampus and prefrontal cortex. IDO1 was normalized GADPH. Data were displayed individually (n =3-4 per group), each sample was repeated 3 times. *** P < 0.005 when compared with control group, ^#^ P < 0.05, ^##^ P < 0.01, ^###^ P < 0.005 when compared with LPS group (one-way ANOVA with LSD test).

### 3.5 ASA VI normalized the aberrant glutamate transmission in hippocampus and prefrontal cortex of LPS-treated mice

To test the effect of neuroinflammatory cytokines and the IDO-mediated kynurenine pathway on neurons, we first examined the morphological and quantitative changes of mature neurons by immunohistochemistry (Fig. 6A). The result showed that the area of NeuN^+^ cell in both hippocampus and PFC were not significant following ASA VI pretreatment regardless of LPS administration as determined by one-way ANOVA (Fig. 6B and 6C). Considering that the effect of the IDO-mediated kynurenine pathway on glutamate transmission may be the main cause of depression-like behavior, we further examined the expression of synaptic proteins (GluA 2 and CamKIIβ). Our analysis of synaptic proteins in mice after LPS administration revealed reduced expression levels of proteins mediating synaptic plasticity CamKIIβ, as well as the AMPA receptor subunits GluA 2 (Figures 6D – 6I). In agreement with the results from the behavioral test, microglial activation, and IDO expression, ASA VI reversed the LPS-induced decrease in the expression of synaptic proteins (GluA 2 and CamKIIβ) in a dose-dependent manner (Figures 6D – 6I). The 40 mg/kg ASA VI consistently normalized the expression of GluA 2 and CamKIIβ both in the hippocampus and the PFC of LPS-treated mice. Although both 10 mg/kg and 20 mg/kg of ASA VI increased the expression of GluA 2 or/and CamKIIβ, the synaptic protein expression levels have not returned to normal. The expressions of GluA 2 in the hippocampus and PFC of LPS-treated mice were normalized by 80 mg/kg ASA VI. But the expression of CamKIIβ only was normalized in the hippocampus of 80 mg/kg ASA VI-treated mice but not in PFC (Figures 6D – 6I).

**Fig. 6.**
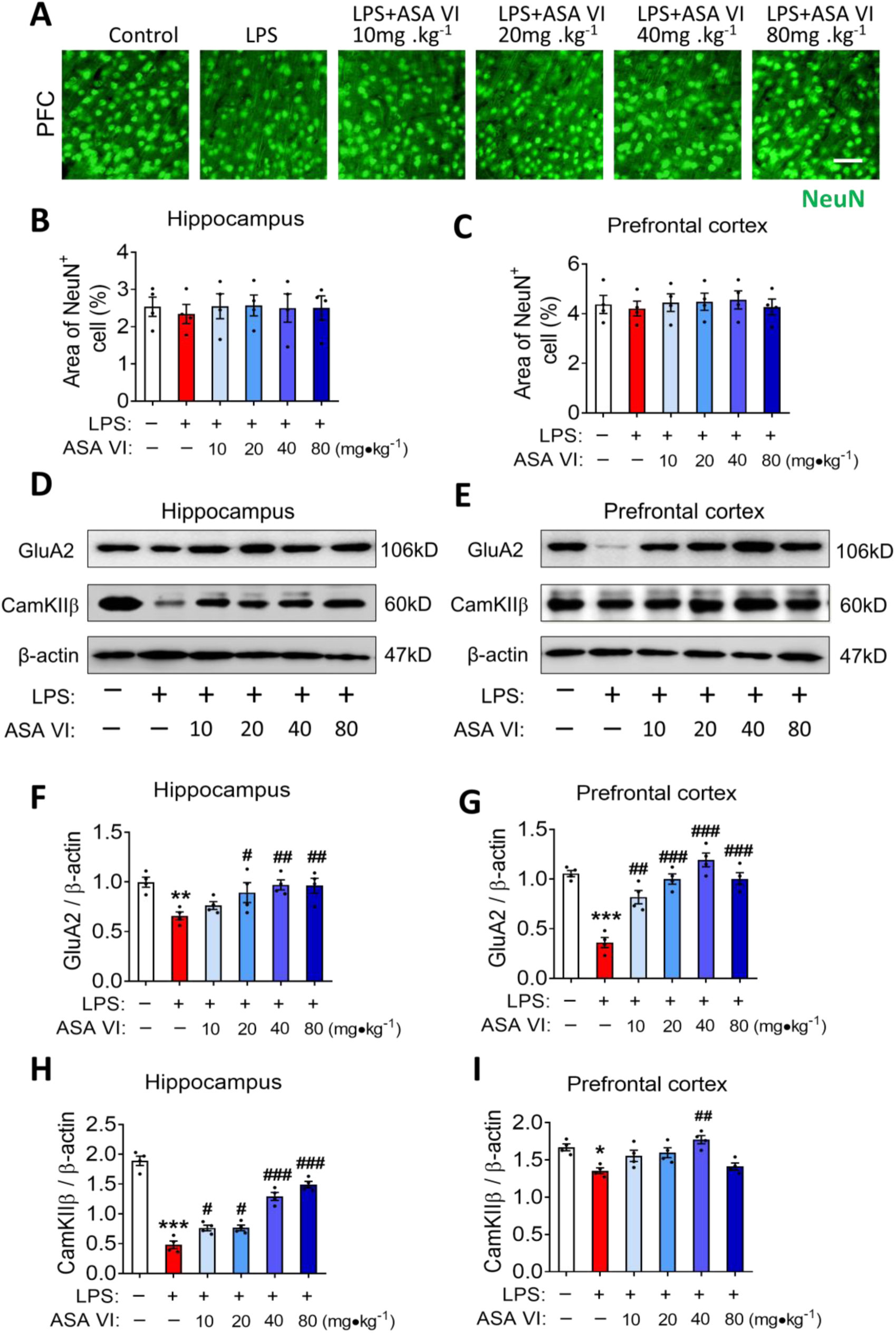
Effects of ASA VI on GluA 2 and CamKIIβ expression in hippocampus and prefrontal cortex of LPS-treated mice. **A**: Immunohistochemistry detects the changes of mature neurons (NeuN, green) in prefrontal cortex of mice following ASA VI pretreatment before LPS or saline administration. Scale bar, 20μm. **B** and **C**: Quantification of the area of NeuN^+^ cells in hippocampus and prefrontal cortex of mice. **D** and **E**: Western blotting detects GluA2 and CamKIIβ in hippocampus and prefrontal cortex of mice following ASA VI pretreatment before LPS administration. **F-I**: Quantification of the protein level of GluA2 and CamKIIβ in hippocampus and prefrontal cortex. GluA2 and CamKIIβ were normalized β-actin. Data were displayed individually (n =3-5 per group), each sample was repeated 3 times for western blotting. 5 immunofluorescence images of each simple were used to analysis. * P < 0.05, ** P < 0.01, *** P < 0.005 when compared with control group, ^#^ P < 0.05, ^##^ P < 0.01, ^###^ P < 0.005 when compared with LPS group (one-way ANOVA with LSD test).

## 4. Discussion

Based on our results, ASA VI showed novel promising antidepressant effects in a validated mouse model of inflammatory depression. Improvements in mice depressive-like behaviors were demonstrated by the reduction of immobility times in the FST test. Furthermore, the mechanism underlying our treatments was investigated by assessing microglial activation, inflammatory expression, NF-κB activation, IDO expression and expression of synaptic proteins. Our results were consistent with previous studies which had indicated that LPS induces microglial activation and IDO expression through upregulation of active pNF-κB [8]. However, the suppressant effects of ASA VI on microglia-mediated neuroinflammatory response, NF-κB activation, and hence the kynurenine pathway are demonstrated for the first time.

ASA VI is the major active ingredient of *Radix Dipsaci*. Although *Radix Dipsaci* is a natural, stable and widely available drug, there are few reports on ASA VI in the literature. There is a small literature showing that ASA VI could cross the blood-brain barrier and has anti-inflammatory and antioxidant effects (Song et al. 2014; Wang et al. 2018; Yang et al. 2019). Based on these, we hypothesized that ASA VI protects the brain from neuroinflammatory damage by inhibiting the expression of microglia-mediated pro-inflammatory cytokines. In this study, the model animals of inflammation-induced depression were successfully established by injection of LPS, which showed the increased in center time in OFT and decreased immobility times in FST. OFT is often used to evaluate the motor ability and anxiety-like behavior of animals(Rajabi et al. 2018). And the latency time and immobility times in TST were usually evaluated the behavioral despair(Yan et al. 2014). We did a dose screening for anti-depressant effects of ASA VI from 10 mg/kg to 80 mg/kg. All of the dose of ASA VI (10 mg/kg 20 mg/kg, 40 mg/kg and 80 mg/kg) did not affect the mobility and stationary time and moved distance both in control mice or LPS-treated mice. These data suggest that ASA VI did not promote excitement or sedation. All of the 20 mg/kg, 40 mg/kg and 80 mg/kg of ASA VI significantly increased the time in center in LPS-treated mice, suggesting ASA VI ameliorated the LPS-induced anxiety-like behavior. The 40 mg/kg and 80 mg/kg of ASA VI significantly reversed LPS-induced increase in immobility times and decrease in latency time in FST, suggesting ASA VI ameliorated the LPS-induced depressive-like behavior. ASA VI did not reserve the LPS-induced decrease in food consumption and body weight. These results suggest that despite the improvement in depression-like behavior, the ASA VI cannot completely eliminate the other effects of LPS injection on animals. Weight and appetite are influenced by many factors, just as LPS affects animals in many ways(Martinez et al. 2014; Subramaniapillai and McIntyre 2017). It is worth noting that 80 mg/kg of ASA VI generally reduced the weight of control mice. The dose may be too high and cause minor damage to the animals.

In the brain, glia could respond positively to LPS stimulation, especially microglia (Zhang et al. 2017). In the study, we found that LPS did not affect GFAP mRNA expression and the area of GFAP^+^ staining. Significantly, LPS administration results in an increase of microglial markers (Iba1 and CD11b) expression and change of the microglial morphology, including the increased area of microglia and decreased number and length of branches in hippocampus and PFC. These results suggest that microglia are more sensitive to immune stimulation than astrocytes(Zhang et al. 2017). When the mice were pretreated with ASA VI before LPS administration, the microglial activation and neuroinflammatory response in the hippocampus and PFC were inhibited by ASA VI in a dose-dependent manner. This result supports our previous hypothesis that the antidepressant effect of ASA VI is potentially mediated by inhibiting LPS-induced activation of microglia and neuroinflammation. Microglia activation plays an important role in the development of depression (Zhang et al. 2018). Neuroinflammatory injury is considered to be one of the main reasons that microglia cells aggravate the pathological process of depression (Zhang et al. 2018). In particular, pro-inflammatory cytokines (such as IL-1β, TNF-α) not only affect the neural function but also induce secondary activation of microglia(Jung et al. 2016; Zhang et al. 2016). ASA VI also inhibited the LPS-induced increase in expression of IL-1β, TNF-α, iNOS, and IL-6 both in the hippocampus and prefrontal cortex.

NF-κB (nuclear factor kappa-light-chain-enhancer of activated B cells) is a protein complex that controls the transcription of DNA (Zhu et al. 2018). NF-κB is found in almost all animal cell types and is involved in cellular responses to stimuli such as stress, cytokines, and bacterial or viral antigens (Pan et al. 2017). NF-κB plays a key role in regulating the immune response to infection(Guo et al. 2019). LPS has been shown to promote microglial activation and neuroinflammatory response by activating NF-κB through their specific receptors (TLR4) (Guo et al. 2019). Our data showed that ASA VI inhibited the LPS-induced phosphorylation of NF-κB in a dose-dependent manner. The results suggest that ASA VI suppressed microglia-mediated neuroinflammation by inhibiting the TLR4/NF-κB signaling pathway (Yu et al. 2012).

Microglial IDO is activated by inflammatory cytokines (including IFN-γ, IL-1β, IL-6, and TNF-α) and psychological stress, facilitate the microglial kynurenine pathway (KP) and result in depressive-like symptoms (Agudelo et al. 2014; Du et al. 2019). The KP was proposed to serve as the switch from acute (sickness-like) effects of inflammatory challenges and stress to the development of depression(Corona et al. 2013). The pro-inflammatory cytokines induced by LPS stimulation can promote the expression of IDO in microglia (O’Connor et al. 2009a). TLR4/NF-κB signaling appears to be the most promising route for IDO elevation in the brain of mice undergoing systemic LPS administration and consequent tryptophan catabolite formations (Hemmati et al. 2019). In this study, we found that ASA VI also inhibited the LPS-induced increase of IDO expression both in the hippocampus and PFC. The results suggest that the antidepressant effects of asperosaponin VI are mediated by the suppression of microglial activation and reduction of TLR4/NF-ĸB induced IDO expression.

Studies in rodents provide even more specific evidence for implicating the activation of the enzyme’s IDO and signaling via the kynurenine pathway in inflammation-induced depression(Wang et al. 2010a; Wang et al. 2010b). The blockade of the KP activation and the neurobehavioral consequences of this activation can be achieved by the IDO inhibitor(Yirmiya et al. 2015). Levels of kynurenine (KYN) and its metabolites 3-hydroxykynurenine (3HK) and kynurenic acid (KYNA) in patients are strongly correlated to depression (Agudelo et al. 2014). The elevated QUIN levels intensely stimulate NMDA receptor signaling, leading to depression(Agudelo et al. 2014). The depressive-like symptoms induced by LPS and other immune challenges were attenuated by NMDA receptor antagonists such as ketamine(Jiang et al. 2019; Walker et al. 2013). Imbalances in glutamate transmission and decreased synaptic plasticity have been suggested as possible mechanisms of depression (Leal and Lopes 2018). Considering that the effect of IDO and inflammatory cytokines on glutamate transmission may be the main cause of depression-like behavior, we further examined the expression of synaptic proteins (GluA 2 and CamKIIβ). Our analysis of synaptic proteins in mice after LPS administration revealed reduced expression levels of proteins mediating synaptic plasticity CamKIIβ and, as well as the AMPA receptor subunits GluA 2. In agreement with the behavioral test results, ASA VI consistently normalized the expression of GluA 2 and CamKIIβ both in the hippocampus and the PFC of LPS-treated mice. The results suggest that ASA VI normalized the aberrant glutamate transmission in the hippocampus and prefrontal cortex of LPS-treated mice.

In conclusion, our results propose a promising antidepressant effect for ASA VI possibly through the downregulation of IDO expression and normalization of the aberrant glutamate transmission. This remedying effect of ASA VI could be attributed to suppress microglia-mediated neuroinflammatory response via inhibiting the TLR4/NF-κB signaling pathway (Fig. 7).

**Fig. 7.**
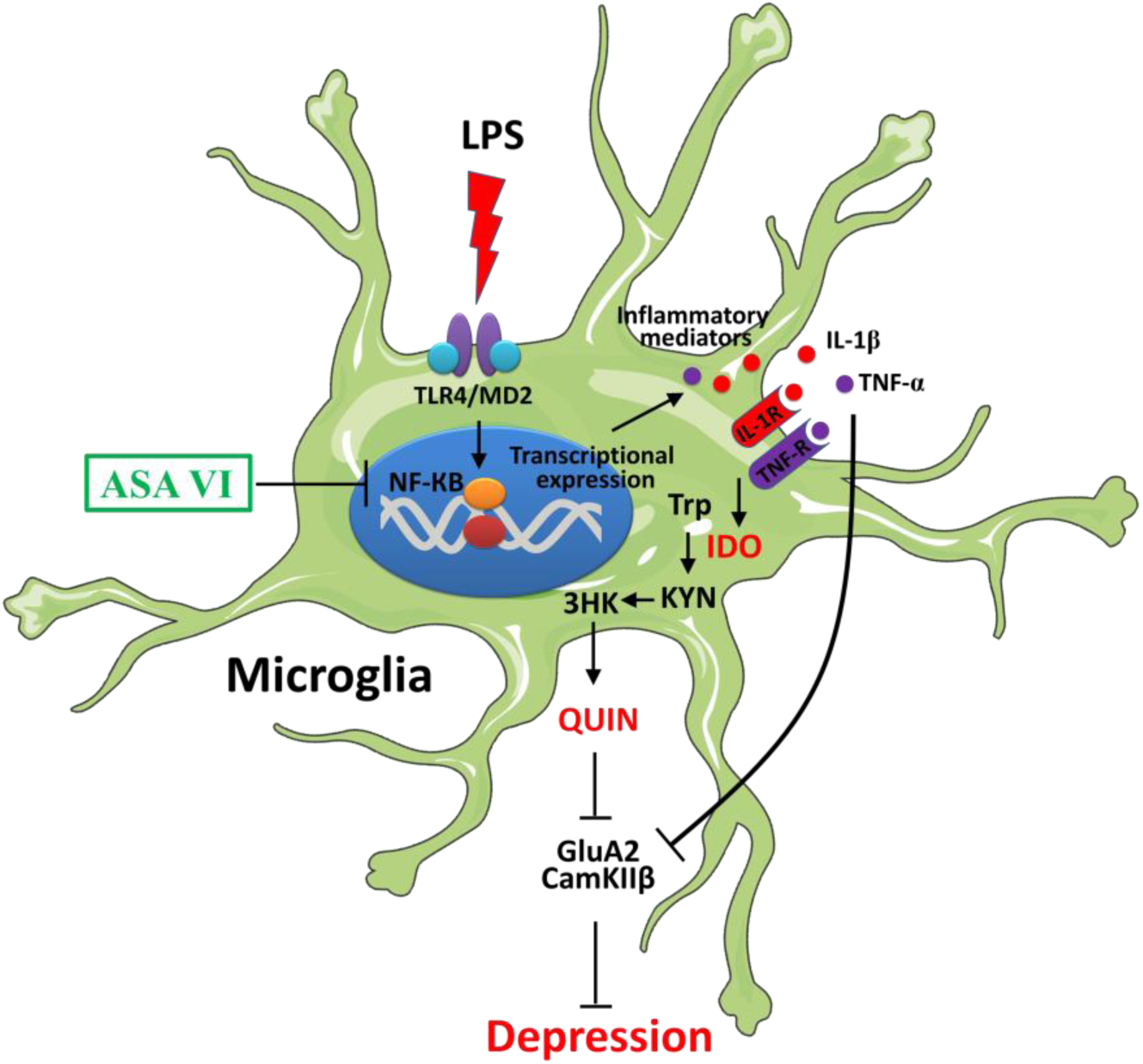
Schematic representation of the mechanism underlying the antidepressant effects of ASA VI. LPS induces the activation of NF-ĸB pathway by TLR4/MD2 in microglia. Activation of NF-ĸB results in the activation of microglia and transcriptional expression of pro-inflammatory cytokines including IL-1β, and TNF-α and IL-6. These cytokines induce the increase of indoleamine 2, 3- dioxygenase (IDO) which catalyze the tryptophan (Trp) into kynurenine (KYN) in microglia. Kynurenine is metabolized to 3-hydroxykynurenine (3HK) in microglia and the 3HK is further metabolized into quinolinic acid (QUIN) which is the agonist of the Nmethyl-D-aspartate (NMDA) receptor that leads to neuronal dysfunction and aberrant glutamate transmission. On the other hand, the pro-inflammatory cytokines also directly leads to neuronal dysfunction and aberrant glutamate transmission. This series of changes eventually leads to depression-like symptoms. Antidepressant effect of ASA VI is potentially mediated by inhibiting LPS-induced activation of microglia and neuroinflammation via inhibiting TLR4/NF-ĸB.

## Acknowledgements

This work was supported by the National Natural Science Foundation of China (81860675), Guizhou science and technology plan project ([2019]5611) and Department of Science and Technology of Guizhou High-level Innovative Talents ([2018]5638).

## Conflict

Authors have no conflict of interest to declare.

## Author Contributions

JQZ, SNY, and TZ designed the conceptual idea for this study and wrote the manuscript. SNY, YHL, CHX,CL performed the experiments and analyzed these data. CGY and WKJ contributed new analytical tools and reagents. All the authors participated in the discussion and approved the manuscript as submitted.

